# Crystallization and preliminary X-ray diffraction studies of melibiose permease

**DOI:** 10.1101/2020.06.26.173740

**Authors:** Yibin Lin

## Abstract

Our work presented here showed that MelB can be crystallized in the conditions as similar as that of other membrane transporter protein of known structure. To identify a rigid protein by modifying the protein structure is the critical factor for facilitating MelB crystallization. It is necessary to perform extensive crystallization screens to obtain crystals. MelB-MelB interaction in the DDM containing solution will be affect by protein preparation, which may lead to reduce in reproducibility of crystallization experiment. Using a detergent mixture is essential for improve protein contact in the crystals, then improve crystallizability. R149C MelB crystal can be obtained in DDM, but these crystals were only diffracted to about 8Å resolution limit. MelB wide type crystal also can be obtained from the condition as that of R149C mutant, but the resolution is weaker than that of mutant. Although MelB and other transporters of known structure share common feature of the crystallization, the emphasis was as much on the protein itself, as it was on detergent type or efficient screening and refinement of the crystallization conditions.

## Introduction

As the highly conformational dynamic of membrane transport protein, it is very difficult to crystallize them and get a good resolution crystal^1-3^. In general, to find a mutant, which may cause a more compact structure and decreased conformational flexibility, is a good way to facilitate the crystallization of this type membrane protein^2,4-6^. For example, lactose permease (LacY), which catalyzes the coupled symport of a galactopyranoside and an H^+^, is a paradigm for the major facilitator superfamily (MFS) of membrane transport proteins^7-9^. LacY wide type couldn’t be crystallized at the beginning^10,11^. However, they found one mutant, the Cys154→Gly mutation causes a more compact structure and decreased conformational flexibility, an alteration that specifically blocks the structural changes necessary for substrate translocation with little or no effect on ligand binding^12-14^. Kaback and his colleagues were successful to crystallize LacY using this mutant^12,15^. In spite of that they had crystallized LacY wide type, but more time and hard work should be need^16^. Clearly, to find a good mutant will facilitate the crystallization.

Furthermore, to find a suit of mutant may help us to understand the mechanism of translocation. Secondary active transport is a form of active transport across a biological membrane in which a transporter protein couples the movement of an ion (typically Na^+^ or H^+^) down its electrochemical gradient to the uphill movement of another molecule or ion against a concentration/electrochemical gradient^17-19^. The model of alternating access, which was put forward more than 40 years ago, is a basic mechanistic explanation for their transport function, and has been supported by numerous kinetic, biochemical and biophysical studies^20,21^. According to this model, the transporter exposes its substrate binding site(s) to one side of the membrane or the other during transport catalysis, requiring a substantial conformational change of the carrier protein^18,22^. Typical over-states include outward-facing, occluded, inward-facing conformation. Clearly, to understand how a transporter protein translocates its substrates through the membrane, it is important to get different over-states structures. Therefore, it is important to crystallize special mutants, which trap protein in special over-state.

Our previous studies showed that R149C mutant preserves the capability of substrates binding, but trap protein in inward-facing conformation^23^. It seems reasonable to suppose that this mutant will decrease the conformational flexibility, and then facilitate the crystallization of MelB. So we decided to use this mutant for crystallization. Previous studies have shown that R141C mutant may display an occlude conformation, in which MelB can bind substrates but cannot translocation^24,25^. So using R141C mutant for crystallization may expect to get a structure, which present an occluded conformation. R149Q may disturb the accessibility of sugar to binding site from extracellular side, but not from the cytoplasmic side, which suggest that R149Q may present a higher possible of an outward-facing conformation^24^. Using this mutant for crystallization may expect to get an outward-facing structure. K377C mutant lost all of the capabilities of substrates binding^26^. Combining all the structures of these mutants, we may explain translocation mechanism of MelB.

## Materials and Methods

### Materials

DDM was obtained from Affymetrix, Inc. Synthesis of the fluorescent sugar analog 2′-(N-dansyl)-aminoethyl-1-thio-d-galactopyranoside (D2G) was carried out by Dr. B. Rousseau and Y. Ambroise (Institut de Biologie et Technologies-Saclay, CEA, France). *E. coli* total lipid extract for the reconstitution of proteins was from Avanti Polar Lipids, Inc. All other materials were of reagent grade and obtained from commercial sources.

### Protein preparation

MelB and the mutants were cloned, expressed and purified similarly as previously reported^27-30^. Follow this method, 8 L of *E*.*coli* cell culture typically produced 15 g of cells, resulting in 5 g of membrane after cell fractionation. From this, 10 mg of MelB R149C mutant could be eluted from a Ni^2+^-NTA affinity column, and 15-20 mg of pure, stable and monomeric R149C was recovered from the size-exclusion column. We observed that the concentration of the detergent used for solubilizing protein from plasma membrane is essential for improving yield. Purified MelB was concentrated using a 50kDa Amicon Ultra-4 (Millipore) and subjected to a buffer exchange process to control the concentration of detergent in protein solution. In this step, about 30% protein will lose.

For protein labeled with seleno-L-methionine, transformed bacteria (B834 DE3) were grown in LeMaster medium (L-methionine replaced by seleno-L-methionine)^31^, and the protein was purified as above with 10 mM DTT added to all buffers. In general, 8 L of cell culture typically produced 10 g of cells. From this, 3 mg of SeMet-R149C MelB could be eluted from a Ni^2+^-NTA affinity.

### Characterization of oligomeric states

Monodispersed, purity, and identity are characterized by gel filtration chromatography during the last step of the purification in an ÄKTA Purifier (GE Healthcare), by Coomassie blue staining SDS-PAGE Gels, silver staining SDS-PAGE gel, and Native PAGE gel.

Gel filtration chromatography separates proteins on the basis of size. Molecules move through a bed of porous beads, diffusing into the beads to greater or lesser degrees. Smaller molecules diffuse further into the pores of the beads and therefore move through the bed more slowly, while larger molecules enter less or not at all and thus move through the bed more quickly. Both molecular weight and three-dimensional shape contribute to the degree of retention. Gel Filtration Chromatography may be used for analysis of molecular size, for separations of components in a mixture, or for salt removal or buffer exchange from a preparation of macromolecules. In this study, size-exclusion chromatography step was used to separate other contaminants. For good separation during size-exclusion FPLC, the protein was applied to the column at ∼5 mg/ml concentration. MelB fractions were collected from the size-exclusion FPLC column at a protein concentration of 0.5-1 mg/ml. In Figure 6.1 we can see the elution profile the gel filtration of MelB from *E. coli* in 0.1 % β-DDM, the symmetry and shape of this profile points to an ideal hydrodynamic behavior for crystallographic studies. Figure 6.2 show coomassie blue staining SDS-PAGE Gel and silver staining SDS-PAGE Gel for the sample, which was ready for crystallization study. The protein purity was estimated at 95-98%.

**Figure 1.**
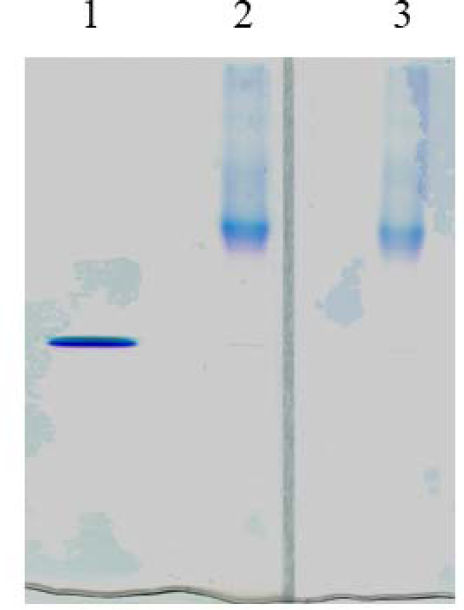
Native PAGE. 1) 10 μ g lysozyme. 2) 10 μ g fresh purified R149C in 0.017% DDM. 3)10 μ g frozen R149C in 0.017% DDM.

**Figure 2.**
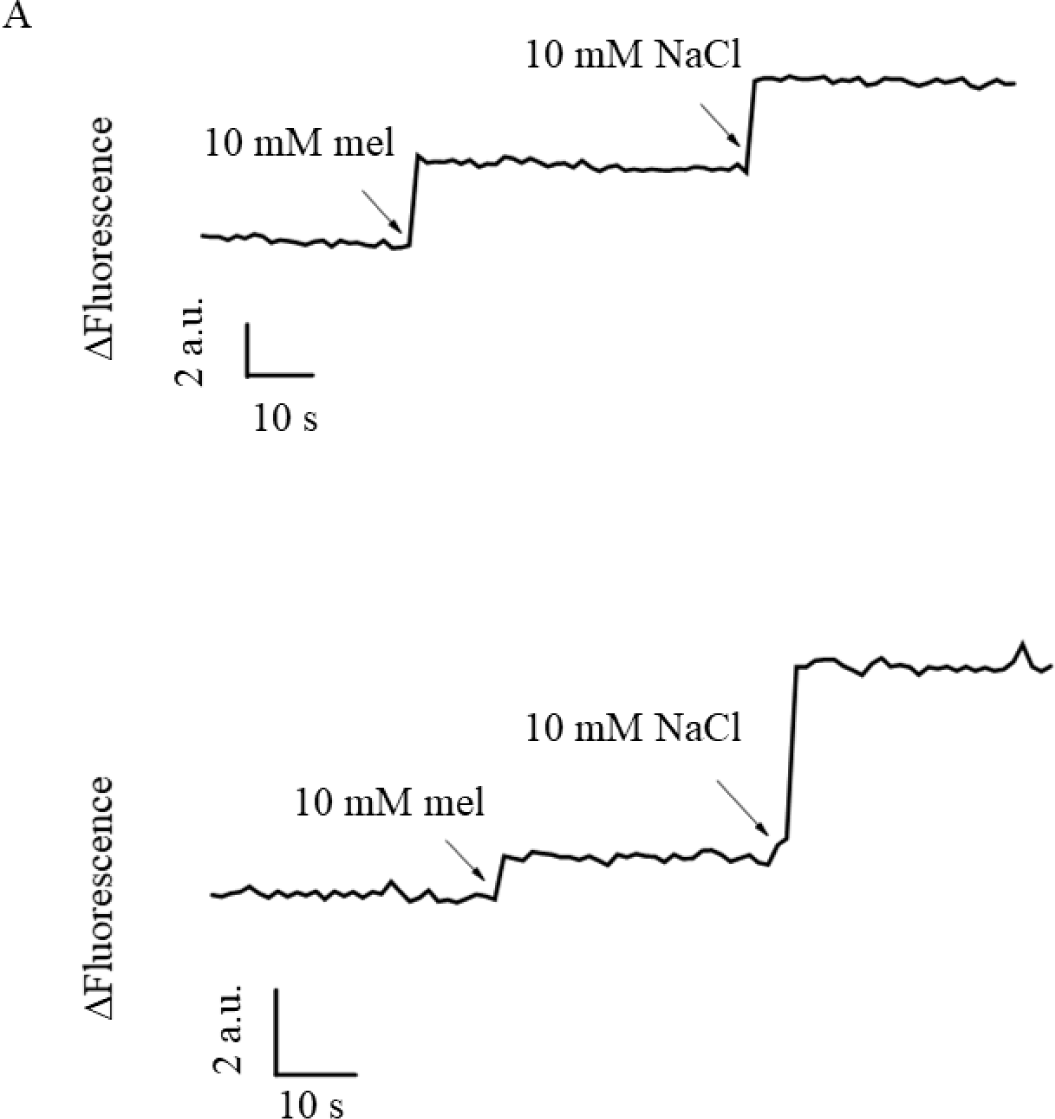
Substrates-dependent Trp fluorescence changes of R149C. (A) Tryptophan fluorescence changes of purified R149C in the buffer containing 0.017% at 20 µg/ml (*λ*_ex_ = 290 nm and *λ*_em_ = 325 nm) after the addition of sugar and Na^+^ to a final concentration of 10 mM (B) Tryptophan fluorescence changes of purified R149C reconsitituted in lipid liposome.

**Figure 3.**
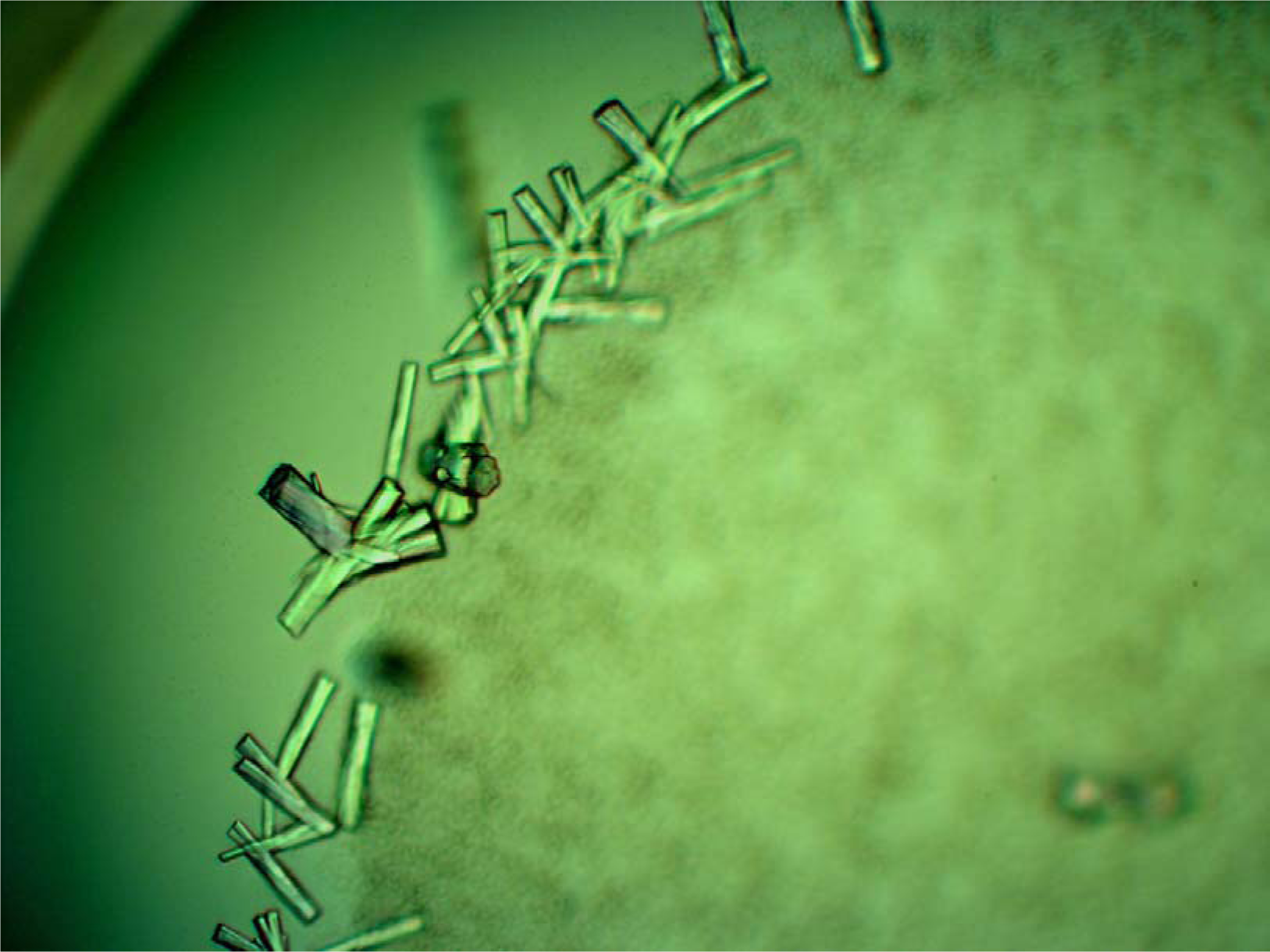
Initial needle-sharp crystals of R149C MelB in β-DDM observed in condition 30 of the JCSG-plus crystal screen.

**Figure 4.**
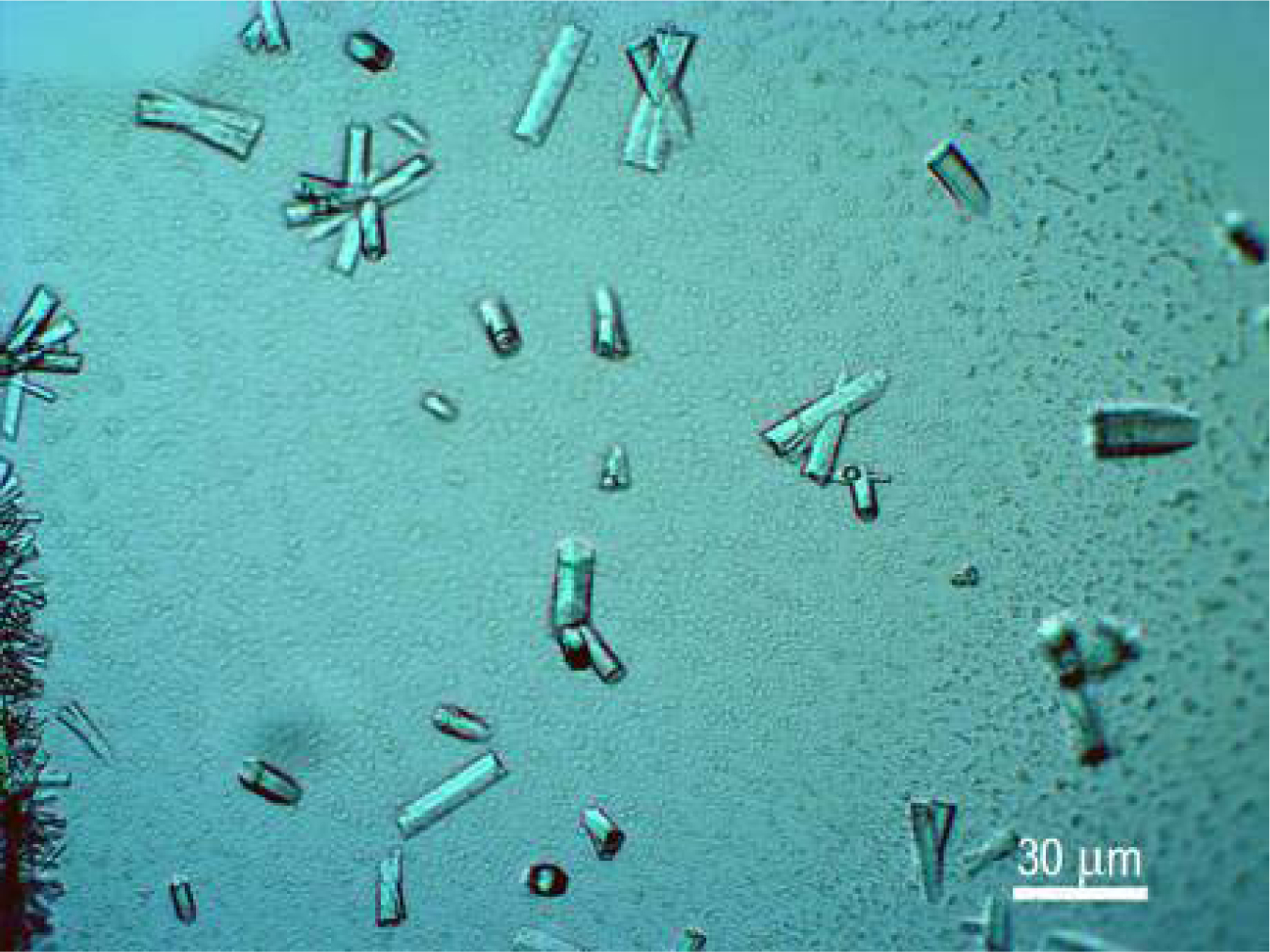
Crystals of R149C mutant form of *E. coli* MelB in the PEG 200 condition.

**Figure 5.**
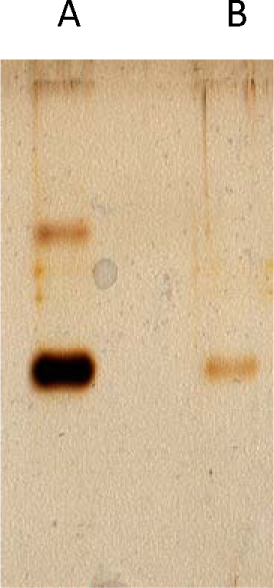
Silver staining SDS-PAGE gel of R149C crystals. A) Crystals; B) control:R149C in detergent containing solution.

**Figure 6.**
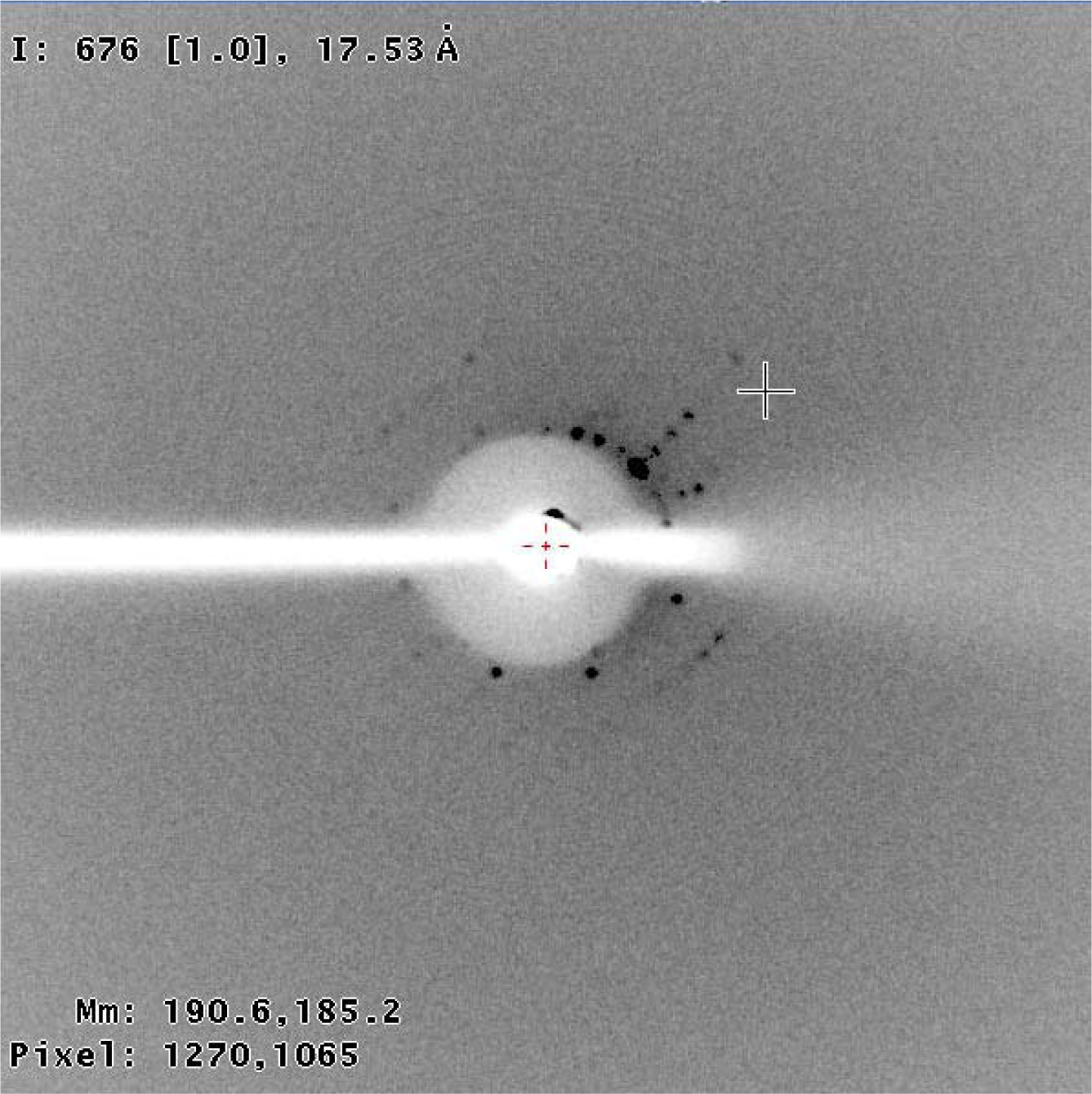
X-ray diffraction pattern obtained using a Cu-Kα radiation from a rotating anode and from crystals similar to those shown in Figure 6. The crystal were flash frozen in liquid nitrogen in its own mother liquor (∼40% PEG 400) which served as cryoprotectant and mounted in a Molecular Dimensions cryoloop/caps. This frame was obtained from a 1 °-1200 s X-ray exposure. The dimensions of this image are 345 mm x 345 mm.

### Crystal preparation

The protein solution was clarified by centrifugation (100,000g, 4 °C, 30 min) before subjecting to crystallization trials. Purified 149C MelB (5-10 mg mg/ml R149C MelB in buffer containing 0.017% β-DDM, 100 mM NaCl, 5% Glycerol, 20 mM Tris-HCl pH 7.5, 2 mM DTT, 10 mM melibiose, and 0.5 mM EDTA) was initially screened using Phoenix RE robot, which sets up the crystallization experiment in 96 well plates using volumes down to 100 nL, against the commercial protein crystal growth screens, e.g., MemGold, MemSys, MemStart, JCSG-plus and PACT (Molecular Dimensions) at both 18 °C and 4 °C. After several attempts of crystallizing the protein during a relatively long time, initial needle-sharp crystals of R149C MelB in β-DDM observed in condition 30 of the JCSG-plus crystal screen at 18°C. We then repeated this condition (changing both the concentration as well as the molecular weight of the PEG used) using the hanging drop vapor diffusion technique by mixing 0.7 μL of protein and 0.7μL reservoir solution at 18°C. Crystals of tetragonal rod appeared from 46-55% (w/v) PEG200 and 29-35% (w/v) PEG400 as precipitant in about one week and grew to full size in about one month. They were frozen directly in liquid nitrogen using PEG as cryoprotectant. We realize that one of the main issues that we had in crystallizing this protein the excess concentration of detergent reached during crystallography sample preparation.

## Results

### Protein preparation

Our studies have shown that when the protein purity is higher than 90%, this should be suit for crystallization studies. However, the homogeneity of the protein is a very important indicator for protein crystallization. Only very high homogeneous protein can be crystallized. Native PAGE electrophoresis is run in non-denaturing conditions. It can be used to determine oligomeric states of the test protein. Figure 1 shows the native PAGE for purified R149C MelB. Only one band was found on this native PAGE, indicating that purified R149C MelB just presents single state. When protein was subjected to freeze-thaw-freeze cycles, the aggregation state of the protein was not changed (Figure 1).

### Trp fluorescence and Trp→*D*^*2*^*G FRET*

To assay the integrity of the R149C MelB in detergent solution, its substrates-dependent tryptophan fluorescence changes upon substrate binding was measured as description in proteoliposome (Figure 2). R149C MelB in detergent solution displays similar substrates-dependent Trp fluorescence changes feature as that of R149C reconstituted in liposome, indicating that R149C MelB in detergent solution preserves the capability of substrates binding and substrates-dependent conformational changes. However, the amplitude of Trp fluorescence changes clearly appears decrease, which may be because of the loose structure in the protein-detergent complex. Then we continue to detect the accessibility of fluorescent sugar analog, D^2^G, to sugar binding site. Unlike in proteoliposome, R149C MelB only displays a few FRET signal with a clear red shift. In agree with the result obtained Trp fluorescence, R149C MelB in detergent solution displays a relatively loose structure. Therefore, R149C MelB in detergent solution preserves the capability of Na^+^ and sugar couple binding.

### The effect of the concentration of detergent

Detergent is quite easily co-concentrated during sample protein preparation when used a Molecular Weight Cutoff Filter, special for that detergents with higher micellar average molecular weight. Too high concentration of detergent will lead to phase separation. Before subjected to crystal screening, purified R149C MelB was concentrated in AmiconUltra 15 centrifugal concentrators (Millipore) with 50 kDa or 100 KDa molecular weight cutoff. We had compared these two type filters, more protein will pass though 100 KDa molecular weight cutoff, which leads the protein lost. Then we went to use 50 KDa molecular weight cutoff. To get an exact concentration of detergent, to avoid co-concentration of any detergent during sample protein preparation, special for those detergents with higher micellar average molecular weight than 50 KDa (for example β-DDM has micellar average molecular weight 50 KDa), the protein sample was buffer exchanged by the batch affinity method by using a small volume of Ni-NTA resin (in a ratio of 15mg protein/1 ml Ni-NTA resin) and eluting by centrifugation the sample in the smallest amount of elution buffer which contained 0.017% β-DDM, 100 mM NaCl, 5% Glycerol, 20 mM Tris-HCl pH8.0, 5 mM DTT, 10 mM melibiose, and 300 mM imidazole (final concentration at 5-10 mg/ml). Then the samples were dialyzed against the crystallization buffer containing 0.017% β-DDM, 100 mM NaCl, 5% Glycerol, 20 mM Tris-HCl pH 7.5, 2 mM DTT, 10 mM melibiose, and 0.5 mM EDTA, for 3h to remove the extra imidazole using a MINI Dialysis Devices (20K MWCO, Thermo SCIENTIFIC). Flow this process, the concentration of the detergent will be control in an expect value.

### Crystal preparation

Here, we recognize that the reason why we did not succeed to crystallize MelB. The main reason is that we suffered from extensive co-concentration of the detergent when concentrating the protein right after the gel filtration procedure. Due to the large dilution effect of the Superdex200 column, we had to concentrate the sample 50-100 x, which in turn also concentrated the detergent to a final concentration of 0.5-2 % from its initial 0.01-0.02 %. All the commercial crystallization screens evaluated with several forms of MelB at this high detergent concentration only yielded spherulite like pseudo-crystalline forms that did not diffract. By using the protein sample containing the nominal lower concentration detergent, we were able to obtain diffracting crystals of R149C MelB. The condition used for crystallization, which is similar to those used to crystallize membrane transport protein is: To form the drops, 1 ul of protein solution in 0.017% β-DDM, 100 mM NaCl, 5% Glycerol, 20 mM Tris-HCl pH 7.5, 2 mM DTT, 10 mM melibiose, and 0.5 mM EDTA at a concentration of about 7-10 mg/ml was mixed with an equal volume of reservoir solution containing 30-50 % (w/v) PEG 200-400 and 100 mM phosphate/citrate pH 4.2 by the hanging drop method at 18 °C. Hexagonal rods appear after 3 days to one week and continue to grow for several more weeks to sizes that range from 10 to 100 μm.

Crystals were washed, dissolved, and identity verified by silver staining SDS-PAGE Gels (Figure 5). Crystals lead a band at same position as that of MelB in detergent containing solution, indicating that the crystals, we obtained in this study was MelB protein. Some bigger crystals were then mounted in cryo-loops and diffracted at 100K in the Plataforma de Cristal.lografia (Parc Científic de Barcelona, Barcelona). Using their rotating Cu anode X-ray source at several exposure times and mar345 image plate detector, we obtained the best macromolecular diffraction to ∼17 Å (Figure 6). The very approximate cell parameters obtained from these images range from 100 - 200 Å which would agree with one molecule of MelB in the asymmetric unit (considering that the MW of the MelB-detergent complex is ∼ 100-120 KDa) and if one considers the following cell parameter/space group combinations: a=b=100 Å c=200 Å in P6 or a=b=200 Å c=100 Å in P622. This crystal was flashed frozen in liquid N_2_ and test at synchrotron (beamline BM16, ESRF) diffracted to about 16Å, which was not higher than that was obtained at home light source.

Clearly, if we intend to gain structural information from the crystals showed in the previous section they need to be improved, that is, their resolution has to be increased from the current 17 Å to at least 4 or 3 Å. Although we expect to improve the crystal diffraction by using synchrotron light, especially when we using microfocus undulator beamline, we are aware that will need careful and possibly lengthy crystal optimization. These methodologies are explained in the following points.

### Crystal optimization by modifying the crystallization conditions

To obtain high quality crystals, screening of a large number condition was essential, special for membrane protein. As Salem Faham and his colleagues reported crystal optimization and structure determination required ∼50,000 crystallization trials and 25 synchrotron trips where more than 2,500 crystals were screened and nearly 120 data sets collected, which shows how difficult is it to get high suit crystals for structure determination^32,33^. A very important feature of membrane protein is that membrane protein is not stable in solution, unless with the help of detergents. However, detergent may interfere with crystal order. It is clear that detergent is the core of membrane protein crystal optimization. Detergent solubilization of proteins entails the formation of protein:detergent complexes or mixed micelles, where easily 50% or more of the total mass of the complex belongs to the detergent. However, historically the interactions between the soluble parts are the first ones to be optimized during protein crystallization. This is a method that has been used for many years in crystallization of soluble proteins. However, such a high percentage of detergent in the protein:detergent complex must have an important effect during crystallization. In addition, the type of detergent is also important to get a stable protein for crystallization. As a matter of fact, it has been observed that many membrane proteins crystallize at detergent concentrations near its cloud point. At this detergent concentration the detergent micelles optimally interact with each other without reaching phase separation. The cloud-point concentration of a specific detergent varies considerably depending on the temperature, pH, precipitant, and concentration of the salts/additives present in solution^34,35^. All of these give us the clues for crystal optimization.

In order to improve the condition that produces the crystal form that we may encounter during this work, the first logical and easiest step is to systematically vary in just a few percent all the variables of the conditions, and setting up a multi-dimensional matrix of conditions in 24-well plates (preferably in hanging drop). Lists of several variables that will have to be modified are:

* Altering the precipitant concentration: changing both the concentration as well as the molecular weight of the PEG used;
* Changing the pH of the crystallization condition: before and after crystallization;
* Varying the drop ratio (protein solution vs. reservoir solution), that is instead of 1:1 μl:μl try 0.5:1, 1:1.5, 0.75:1, 1:0.75, etc.;
* Testing different temperatures;
* Modifying the protein concentration used for crystallization (variations of 20% above and below the current protein concentration);
* Extensive screening for small molecule additives;
* Trying to extract protein from membrane with different detergents;
* Trying to purify protein with different detergents;

### Effect of the precipitants

Figure 7 summarized the different precipitants and the concentrations of the polyethylene glycols (PEGs) used in the successful crystallization of the membrane proteins. Unlike soluble proteins, small MW (molecular weight) PEGs, in particular PEG 400, have been more successful for membrane proteins (Figure 7A). Figure 7B shows the optimum concentration range for low MW PEGs lies between 20% and 30%, which is relatively narrow compared to the large MW PEGs. As other transporters, we obtained the MeB crystals from the condition using PEG200-400 as precipitant. To find a suit precipitant for R149C MelB crystallization, we fixed buffer, change the MW and the concentration of PEG. We tested PEG200 at 30-60%, PEG300 at 20-50%, PEG400 at 10-40%, and PEG500 at 5-25%. Crystals appear from PEG200 at 46-55%, PEG300 at 40-50%, and PEG400 at narrow range, i.e., 29-35% (Figure 8). No crystal was obtained from the PEG with MW higher than 400. These results show that only small molecular weight PEGs are suit for MelB crystallization. In contrast, we tested some traditional precipitants, e.g. (NH_4_)_2_SO_4_, which were used for protein crystallization. Only brown precipitation was obtained at any concentrations, which indicates that only low MW PEGs are suit for the stable of MelB-DDM complex. The best crystal was diffracted to 11Å at ESRF BM16.

**Figure 7.**
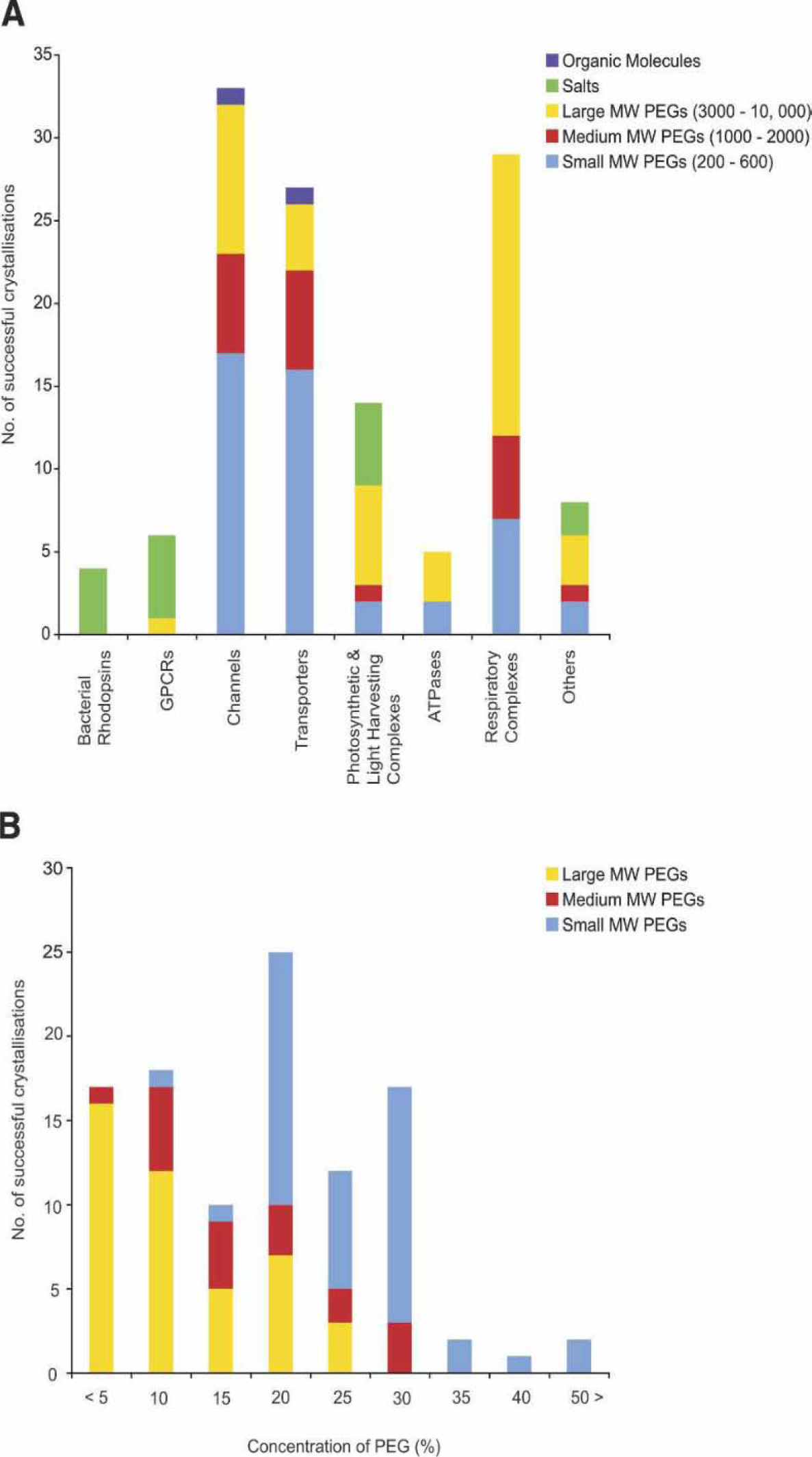
(A) The different precipitants used in the successful crystallization of the membrane protein families are shown. (A) Precipitants. (B) The concentrations of the polyethylene glycols used for successful crystallization of the membrane protein families are shown. (according to Newstead et al. 2008)

**Figure 8.**
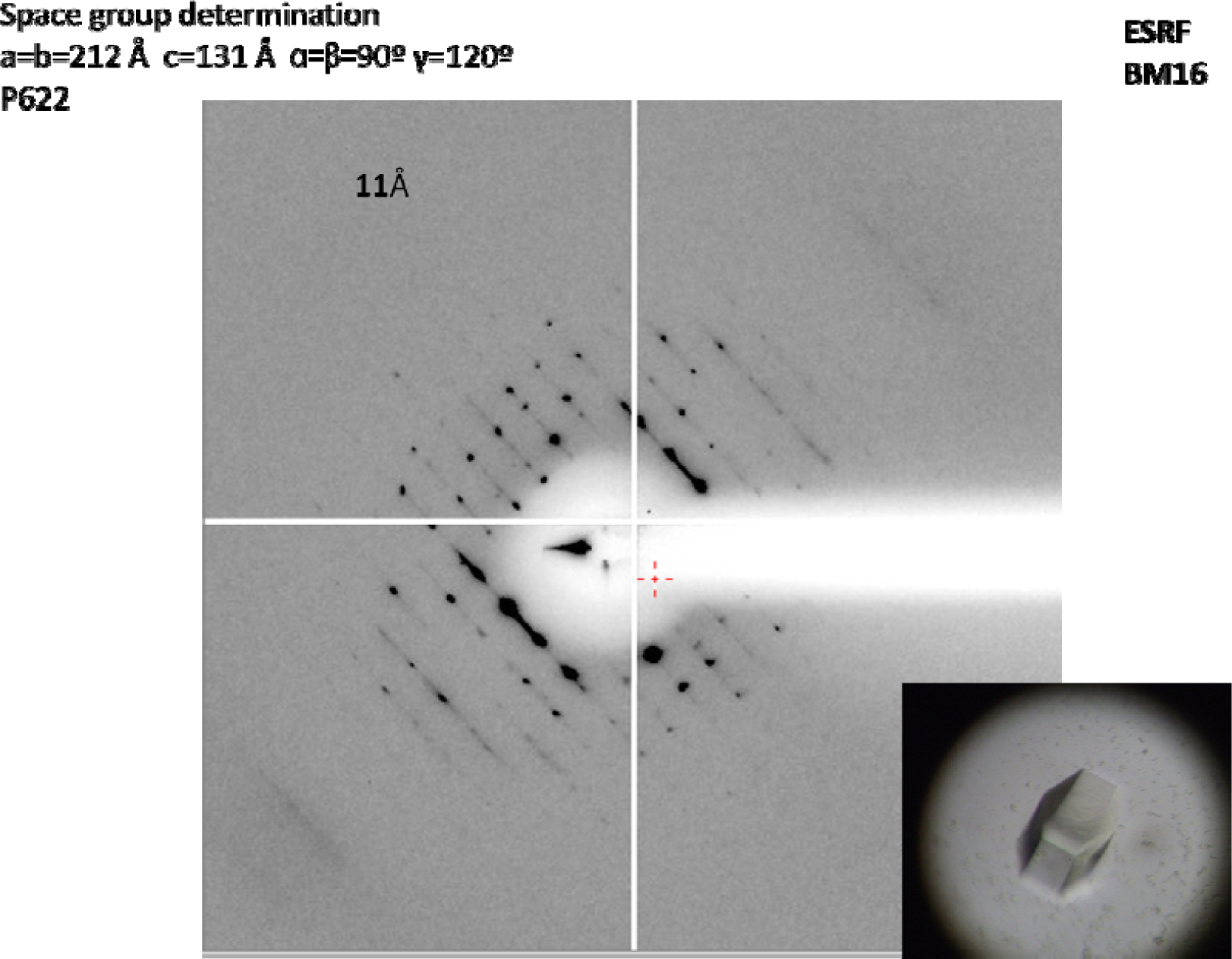
MelB crystal and X-ray diffract pattern.

### Effect of the pH in the reservoir solution

pH is an effective crystallization variable because most proteins demonstrate pH dependent solubility minima and will solubilize, precipitate, or crystallize at particular pH values. To test the effect of the pH in MelB crystallization, we fixed precipitant, i.e., 48% PEG200, and then varied the pH by 0.1 pH unit from 3.5 to 9.0. Similar rod-like crystals were obtained sodium citrate buffer at pH 4.0 to 4.5. This buffer change neither clearly improves the size of the crystals nor improves the resolution of the crystals. Only need-like crystals were obtained from MES buffer with pH at 6.5. Some bigger crystals were only diffracted to 18-20 Å at synchrotron (ESRF BM16 beamline). The study of buffer screening suggested that phosphate/citrate pH 4.2 may be the best buffer for MelB crystallization.

### The effect of the concentration of protein

Initially, we obtained R149C MelB crystals at the protein concentration of 9.2 mg/ml. Then we test series of the concentration of protein from 4 mg/ml to 11 mg/ml by fixing buffer (phosphate/citrate pH 4.2) and precipitant (48% PEG200) in hanging drops at 18 °C by equilibrating a 1:1 mixture of protein and reservoir solutions against the reservoir. It seems that R149C MelB perfects to be crystallized at the concentration from 4.5 to 9 mg/ml. Another easy way to test the effect of the concentration of protein is by varying the drop ratio (protein solution vs. reservoir solution), that is instead of 1:1 μl:μl try 0.5:1, 1:1.5, 0.75:1, 1:0.75, etc. The result showed that using the ratio of 0.7μl protein to 1.4μl reservoir solution will facilitate the R149C MelB crystallization. At this ratio, crystals will appear in 3-7 days.

In contrast, at 1:1 ratio crystals will appear in more time, typical appearing in two weeks. However, 1:1 ratio will be much easy to obtain large crystals and less crystal in one drop. Although, 1:2 ratio is much easy to obtain crystals, crystals also are small and there are many crystals in one drop.

### The effect of the substrates

It is clear that to get a stable protein is essential for crystallization and getting high quality crystals. Membrane protein is characterized by its highly dynamic. The binding of substrates may help to stable protein. However, some persons have reported that higher substrates concentrations in the drop lead the decrease of the resolution^36^. For MelB, there are two substrates, i.e., Na^+^ or Li^+^ and melibiose. Na^+^ is one of the substrates, but it is also play an important role as salt in crystallization. From the already reported structures of this type transporter, which couple transport substrates with sodium, they almost used NaCl in 50 mM to 100 mM. And the organic substrates, e.g., sugar, amino acid, nucleic acid, etc. were used in a lower concentration, e.g., 2-5 mM, always closing to their binding constants. To test the effect of the substrates, we tried to change the concentration of substrates in the protein solution or added them to the reservoir solution as additives. To change the concentration of substrates in the protein solution, we modified the dialysis buffer by changing the concentration of Na^+^ or melibiose. In this study, we test different concentration of NaCl from 20 mM to 150 mM and melibiose from 2 mM to 20 mM melibiose. The results show that crystals appear from all of these conditions. However, higher concentration of the substrates is easy to obtain small, but many crystals in drop. Lower concentration of the substrates is much difficult to obtained crystals under the same condition. So finally we had chosen 100 mM NaCl and 10 mM melibiose for R149C MelB crystallization.

### The effect of temperature

From the already reported structures of this type transporter, almost proteins were successfully crystallized at 18-20°C or 4°C. In this study, we test two different temperatures, i.e., 18°C and 4°C under the same condition. The results show that R149C MelB perfects to be crystallized at 18-20°C. We also obtained crystals at 4°C. However, crystals grew slower than that at higher temperature (more than one month). Another thing is that crystals growing at 4°C are always small. When we diffracted these crystals obtained at 4 °C, there are not any crystals which can diffract to lower than 17 Å at synchrotron (ESRF ID 23-1 beamline). These results suggest that lower temperature may not suit for the crystallization of R149C MelB. Then we chose 18 °C for further crystal optimization.

### Effect of phospholipids (PL)

Phospholipids can also have a positive impact in the crystallization of membrane proteins solubilized with detergents. Of the transporters that are most similar to MelB, two of them: *E. coli* GlpT and LacY require phospholipids for its crystallization or their presence during crystallization increased significantly the quality of the obtained crystals^37,38^. During the purification of GlpT, it was observed that endogenous phospholipids co-purified with the protein in molar ratios that ranged from 1:20 to 1:40 (protein:phospholipid)^39^. Guan and her colleagues reported three different crystal forms that diffract to increasingly better resolution in a manner that correlates with the concentration of copurified phospholipids, i.e., mol PL/ mol LacY 8→5Å; mol PL/ mol LacY 18-25→3Å; mol PL/ mol LacY 9-16→2.6Å. In the case of LacY, and similar to GlpT, the addition of *E. coli* phospholipids to purified protein was a requirement to obtain crystals of sufficient quality of the wild-type protein.

To test the effect of the phospholipids in MelB crystallization, we used the same method employed in the cited work. To this end, we tried to directly add *E*.*coli* phospholipids (Avanti Polar Lipids, Inc.) to reservoir solution as additive or tried to purify protein with the addition of *E*.*coli* phospholipids. In this study, we test the effect of phospholipids with the protein in molar ratios that ranged from 1:5 to 1:40 (protein:phospholipid). It seems that the addition of phospholipids in reservoir resolution may not help to facilitate MelB crystallization. The problem is phospholipids cannot solve in water, but in detergent containing buffer. In general, phospholipids were solved in 0.5% β-DDM. However, in the reservoir solution, there is not detergent or just 0.017% (double to CMC). When we added phospholipids to reservoir solution, almost lipids precipitated. Then we went to purify protein with the addition of phospholipids to the washing buffer and eluting buffer. Our results showed that with the help of lipids, the crystals were grown within one month to optimal size (800×60×60 μm^3^). The best crystal was diffracted to about 8 Å at synchrotron ESRF ID 23-1 beamline (Figure 9A) and about 9 Å at synchrotron ESRF ID 14-4 beamline (Figure 9B).

**Figure 9.**
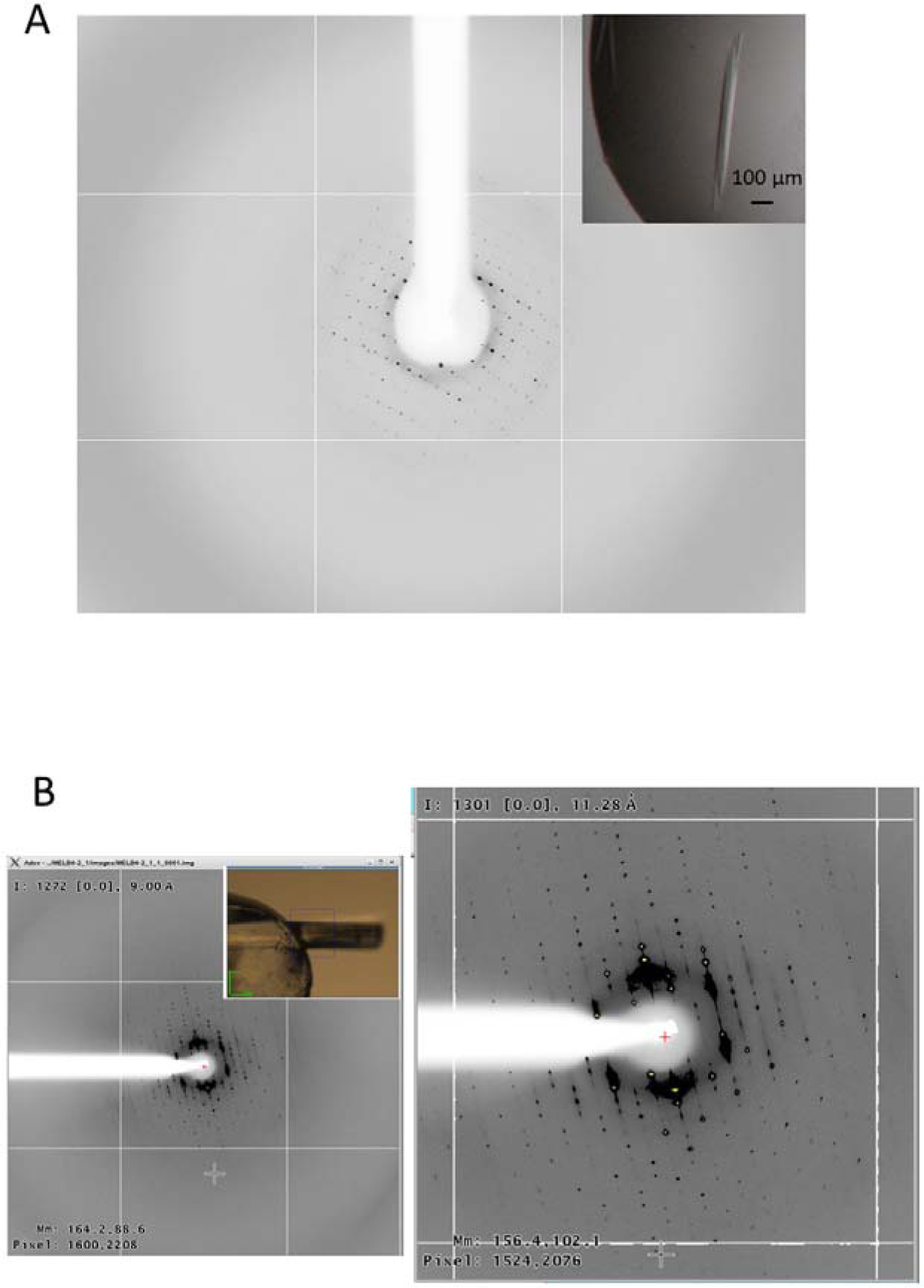
X-ray diffraction pattern obtained at synchrotron ESRF ID 23-1 beamline (A) and at synchrotron ESRF ID 14-4 beamline (B). The crystals were flash frozen in liquid nitrogen in its own mother liquor (∼48% PEG 200) which served as cryoprotectant and mounted in a Molecular Dimensions cryoloop/caps. Insert figure in A shows the photo of crystal using for X-ray diffraction study. (B, right) shows enlarged image of (B, left) to show detail diffraction spots. Insert figure in B shows the photo of crystal which was mounted on the cryoloop.

## Discussion

Inspired by the successful structure determination of about 40 membrane transport protein, we took a similar approach to crystallize MelB. The crystallization of the MelB is summarized in Figure 6.23. The key elements in our experiments were to i) indentify a rigid protein core by modifying the protein construct; ii) prescreen detergents for their ability of maintaining protein monodispersity; iii) perform extensive crystallization screens for every detergent identified in the previous step and repeat any conditions in several times to improve reproducibility; iv) improve protein contact in the crystals by using a detergent mixture. Hence, the emphasis was as much on the protein itself, as it was on detergent type or efficient screening and refinement of the crystallization conditions.

Secondary transporter uses energy to transport molecules across a membrane by series of conformational changes. In general, the translocation is expected to take place by means of an alternating access mechanism, where the structurally coupled substrates binding sites present access to the extracellular medium in the binding process and to the cytoplasmatic medium in the release process^40^. This mechanism has been supported by numerous kinetic, biochemical, biophysical studies and atomic-resolution structural evidences recently^41^. The biggest characteristic of these proteins is dynamic, needing for translocation of substrates, but making them difficult to crystallize. In many cases, finding a grid protein core by muting or trimming original protein will facilitate the crystallization and help to improve the resolution. For example, lactose permease was firstly crystallized using a mutant, i.e., C154G, which blocks protein in inward-facing conformation. Higher resolution crystals of GlpT were obtained by trimming original protein. In our case, we attempted several times to crystallize MelB wide-type, but not successful. The breakthrough was that we found one mutant, i.e., R149C, which can bind substrates, but fixes MelB in single conformation. We were successful to obtained R149C MelB crystals from some conditions as that of other secondary transporters. We summarized R149C MelB crystallization experiences, and developed a simplified grid screen. MelB wide type was then subjected to this grid screen. We obtained MelB crystals from some condition as that of R149C MelB. In contrast, no crystal was obtained from R149Q MelB, R141C, and K377C MelB when we performed same grid screen. These interesting results indicate MelB wide type and R149C MelB should share something similar structure feature. In agreement with the studies of crystallization, previous studies, based on analysis of infrared spectroscopy and fluorescence spectroscopy have shown that all of these three mutants disorder MelB native structure in some degree. Although, MelB crystals could be obtained from same conditions at that of R149C MelB, the resolution is clearly lower than that of R149C MelB, which may imply the difference of protein-protein interaction in crystal pack. One reasonable assumption may be that R149C mutant fixes MelB in a relatively stable conformation, which reduces the conformational dynamic, then is benefit to form a stable crystal cell. Our previous studies as described in PART I, showed that R149C MelB trapped MelB in inward-facing conformation. In short, to indentify a stable structure that reduces the conformational dynamic is essential necessary for obtaining higher resolution crystal.

We observed a critical factor for obtained membrane protein crystals is the concentration of detergent in protein solution. Due to the high hydrophobicity, membrane protein should be prepared with the help of detergent. However, excess detergent will disturb the formation of protein crystals, which may be due to reduce protein-protein interaction, and then reduce crystallizability. In general, protein crystallization depends on random protein-protein interactions^42^. Membrane protein will be surrounded by excess detergent micelles, therefore exposes less protein surface areas for protein-protein contacts. On the other hand, too much detergent can denature the protein or impede crystallization by phase separation^43^. We observed that the main reason, which causes excess detergent, is that detergent is co-concentrated during sample protein preparation by a means of Molecular Weight Cutoff Filter. For MelB, this effect should be quite important. None crystal was obtained only when we well control the concentration of detergent. We had tried to use 50 kDa or 100 KDa molecular weight cutoff to concentrate protein sample. In any cases, none crystal was obtained, implying either 50 KDa or 100 KDa molecular weight cutoff co-concentrated detergent during protein preparation. Actually, many persons reported that they used 50 kDa or 100 KDa molecular weight cutoff to concentrate protein in the presence of DDM ^44^. Therefore, MelB should be quite sensitive to the concentration of detergent. We may suppose that the protein interactions of MelB in DDM containing solution should be very weak, which may be the cause of the difficulty of the crystallization of MelB. Another case, MelB seems quite stable in DM and B-NM, can be concentrated to higher concentration in these two detergents. However, when we performed crystal screen for these samples, none crystal was obtained, which suggests that there is only detergent-mediated interactions. Our observation implies DDM, DM, NM, which significantly protect hydrophobic protein may be suit for the stability of MelB, but this type detergent reduces protein-protein interactions, and then reduces crystallizability. Mixture detergent method may be effective to improve crystallizability and resolution.

Extensive crystallization screens were essential to obtained crystals. Sparse matrix screens allow sampling of a broad range of conditions screens that are more systematic with respect to pH as well as precipitant type and concentration can be used. We used a number of commercially available screens: MemGold, MemSys&MemStart, JCSG-plus and PACT (Molecular Dimensions). MemGold designed for membrane proteins studied, is based on the conditions that have successfully generated a-helical membrane protein crystals used to solve X-ray structures^45^ and contains 96 conditions covering a range of pH, PEGs and salt additives. MemStart is a sparse matrix screen, whereas MemSys is a systematic exploration of pH, salt concentration/type and precipitant concentration/type. MemSys&MemStart was designed for screening and optimizing crystallization conditions for transmembrane proteins using vapour diffusion methods. JCSG-plus is a 96 reagent, optimized sparse-matrix screen of classic and modern conditions. PACT has been developed to systematically test the effect of pH, anions and cations, using PEG as the precipitant.

A survey of the published literature reveals that small MW PEGs, in particular PEG 400 have been more successful for a-helical membrane protein crystallization, as the precipitant. Organic solvents tend to disturb the detergent micelles and, at high concentrations, denature membrane proteins. Salt, on the other hand, reduces the solubility of the detergent micelles and often precipitates the membrane protein embedded in the detergent micelle before crystallization occurs. For MelB, crystals were obtained from the conditions using small MW PEGs as precipitant. PEGs are therefore a best choice for membrane protein crystallization. Another important factor is pH. More than 90% of soluble protein crystals are grown between pH 4 and 9. We surveyed the entire buffer which was used for the membrane transporter protein of known structure, suggesting this will be true for membrane proteins as well. Thus, we used screening kits with 4 small MW PEGs over the pH range of 3.5 to 9.0 based on the initial crystallization results of the protein in DDM. Indeed, their application to MelB purified in several detergents produced numerous hits, which were subsequently pursued for optimization. Furthermore, the utilization of these screening kits significantly improved the efficiency and reproducibility of crystallization experiments, which was particularly beneficial when a dozen of detergents were screened for crystallization. Finally, the 4 PEG/pH screening kits used to search for nucleation of R149C MelB crystals may serve as a starting point for other mutant form MelB.

The membrane protein molecule in solution exists as a protein-detergent-lipid complex. Specific lattice contacts in any protein crystal are made exclusively via protein-protein interactions, and too large a detergent micelle can be an obstacle for protein crystallization. By reducing the heterogeneity of the protein surface and by optimizing the detergent micellar size and shape, we essentially increased the area available for the formation of lattice contacts, thereby improved the protein crystallizatbility. In many cases, second detergent or third detergent has shown to be essential for improving the diffracting resolution. For example, Wang, Y. and his colleagues reported that for formate transporter FocA, Cymal-2 was shown to be essential for improving the diffracting resolution from 3.5 Å to 2.2 Å. Another case, Lemieux, M. J. and her colleagues reported that a detergent mixture of DDM and C12E9 is requirement to give crystals that diffracted to 3.3 Å resolution, but DDM alone resulted in crystalline order to 7 Å, complete detergent exchange to C12E9 gave no crystals^39^. Based on the data obtained from the transporters of known structure, we found β-DDM, β-UDM, β-DM, β-OG, and lipid-like detergents, e.g., Cysmal-6 were quite common used to extract protein from plasma membrane. And C12E8, LDAO might use as second detergent for protein preparation. In order to improve efficiency, simplify the experiment, combined with published experimental data and our experimental data of R149C MelB, we have listed some of the most commonly used detergents i.e., β-UDM, β-DM, LDAO, β-OG, β-NM, β-NG, β-OM, α-DDM, C8E4, C12E8, C12E9, Cymal-5, Cymal-6, Fos-choline-12. Then we went to prepare MelB or other mutant forms MelB with β-DDM, β-UDM, β-DM, β-OG, and Cymal-6 and performed detergent screen as districted at up sentence. Following this method, we were successful to obtained MelB wide type crystal, which was never crystallized, and many different sharp crystals of R149C MelB or MelB were obtained by this screening.

In our studies, we note that unlike soluble protein, although we are strictly in accordance with the same experimental procedures, it is quite difficult to reproduce MelB crystals. We believe that the problem lies in the protein preparation. In fact, our observation showed that only a small percentage of new preparation protein can be crystallized. It may be due to the unstable of protein-detergent complex (PDC). It is clear detergent micelle is not as stable as lipid liposome or plasma membrane. Temperature, mechanical force, etc. would affect the stability of PDC, then disorder protein conformation. There may be two possibilities: i) protein-detergent complex is disordered, detergent cannot protect hold protein. In this case, membrane protein will precipitate; ii) protein-detergent complex is disordered, detergent will protect hold protein, none protein surface will expose to outside. In this case, there is only detergent-mediated protein-protein interaction, which will reduce the crystallizability. Clearly, the reason resulted in lower reproducibility, should be the latter case. In fact, it is inevitable to change the surface of the protein. The biggest difficulty is that there is still not effective method used to detect the types of protein-protein interaction. So we should perform extensive crystallization trials for purified proteins. To improve the efficiency and reproducibility of crystallization experiments, it is necessary to establish a high-speed and convenient crystal screening methods. Based on the data obtained from some of other transporters of known structure and the experience of the crystallization of MelB, we developed a simple crystal screening method combining sparse matrix screen, detergent screen, and additive screen.

MelB crystals were obtained from the conditions as similar as that of other transporters of known structure. This interesting phenomenon implies that the crystallization of such proteins may follow the common law. In agreement with this phenomenon, all secondary transporters share some common structural features, e.g., transmembrane α-helix secondary structure, heart-shaped with an internal cavity and an approximate two-fold symmetry, higher conformational dynamic, etc, which determine all of these type protein should follow the common law. For example, almost transporters were crystallized using PEG400 as precipitant. The protein buffer and crystallization buffer for different proteins are quite similar. Almost crystals were obtained at 18-20°C. The difference of crystallization conditions between different proteins may be because of the difference of surface property, dimension, stability, etc. These transporters are hydrophobic and dynamic, making them difficult to crystallize. The core work of the crystallization of this type protein is to find a rigid protein to reduce dynamic and a suit detergent or detergents mixing to improve the stability of the protein and the protein-protein interactions in the crystal pack. Mutagenesis is an extremely popular and useful method to prepare structure-specific proteins. In many cases, more stable structure of the protein can be obtained by site-direct mutating or trimming original protein. As the difference of the dimension, surface property, the number of transmembrane α-helixes, to find a suit detergent for specific membrane protein crystallization, large detergents should be screened either for protein preparation or crystal screens. Size exclusion chromatography, infrared spectroscopy and fluorescence spectroscopy have been used in detecting the stability of the modified protein or detergent-protein complex. Another feature of membrane protein crystallization is very low reproducibility. As the detergent micelle should not be as strong as plasma membrane or lipids liposome, the stability of membrane protein structure will be strongly affected by temperature, mechanical force, and other physical and chemical constants. All of these factors would lead that every time we got the protein which was not completely consistent, although we were strictly in accordance with the same method. So repeated in several times is essential need to obtain crystals and improve resolution.

---

**Figure 12.**
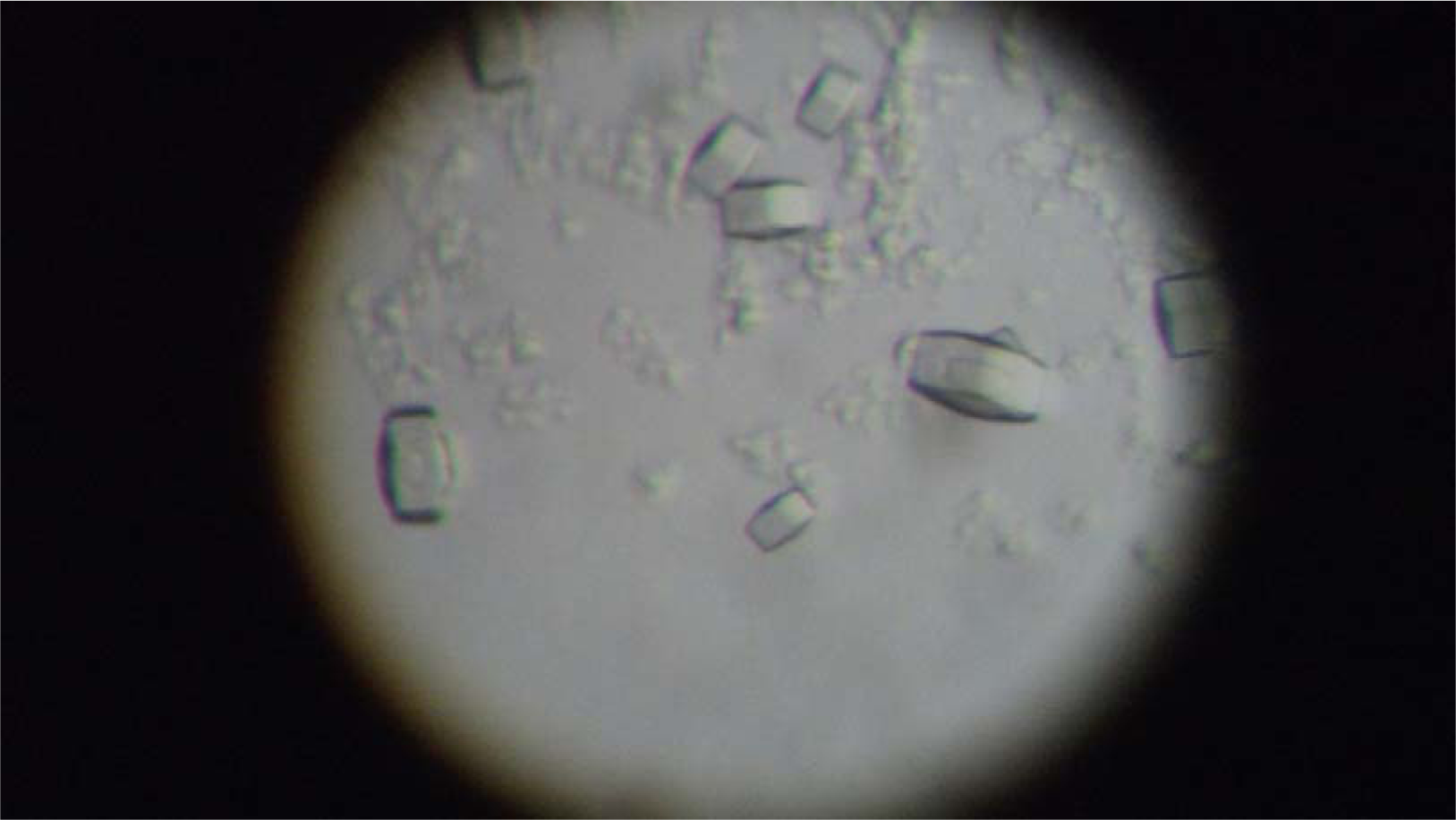
crystals growing at 4 degree.

